# An ensemble method for designing phage-based therapy against bacterial infections

**DOI:** 10.1101/2022.06.01.494305

**Authors:** Suchet Aggarwal, Anjali Dhall, Sumeet Patiyal, Shubham Choudhury, Akanksha Arora, Gajendra P.S. Raghava

## Abstract

Phage therapy is a viable alternative to antibiotics for treating microbial infections, particularly managing drug-resistant strains of bacteria. One of the major challenges in designing phage based therapy is to identify the most appropriate phage to treat a bacterial infection. In this study, an attempt has been made to predict phage-host interaction with high accuracy to identify the best virus for treating a bacterial infection. All models have been developed on a training dataset containing 826 phage host-interactions, whereas models have been evaluated on a validation dataset comprising 1201 phage-host interactions. Firstly, alignment based models have been developed using similarity between phage-phage (BLAST_Phage_), host-host (BLAST_Host_) and phage-CRISPR (CRISPR_Pred_) where we achieved accuracy between 42.4%-66.2% for BLAST_Phage_, 55%-78.4% for BLAST_Host_, and 43.7%-80.2% for CRISPR_Pred_ at five taxonomic levels. Secondly, alignment free models have been developed using machine learning techniques. Thirdly, hybrid models have been developed by integrating alignment-free models and similarity-score where we achieved maximum performance of (60.6%-93.5%). Finally, an ensemble model has been developed that combines hybrid and alignment based model. Our ensemble model achieved highest accuracy of 67.9%, 80.6%, 85.5%, 90%, 93.5% at Genus, Family, Order, Class and Phylum levels, which is better than existing methods. In order to serve the scientific community we have developed a webserver named PhageTB and standalone software package (https://webs.iiitd.edu.in/raghava/phagetb/).

**Key Points:**

- Phage therapy provides an alternative to mange drug resistant strains of bacteria
- Prediction bacterial strains that can be treated by a given phage
- Alignment-based, alignment-free and ensemble models have been developed.
- Prediction of appropriate phage/virus that can lyse a given strain of bacteria.
- Webserver and standalone package provided to predict phage-host interactions.

## Introduction

Bacterial infections pose a major threat to public health across the globe. According to recent reports around 1.27 million people died in 2019 by bacterial infection due to antimicrobialresistance [1]. In the last few decades, heavy consumption and misuse of antimicrobial and antibacterial drugs have exacerbated the current crisis [2, 3]. It has been observed in recent studies that number of novel bacterial strains are emerging which are resistant to existing antibiotics [4]. Therefore, researchers are looking for alternative approaches, one such approach is “phage therapy” where viruses infect and lyse bacterial strains [5-8]. One of the major challenges in designing phage therapy is to identify the most efficient virus or bacteriophage that can lyse a target strain of bacteria [9, 10]. Currently, several traditional techniques such as RNA-sequencing, microfluidic-PCR, PhageFISH, flow cytometry have been used to measure the phage-host interactions. Though these experimental techniques are highly accurate in identification of virus-bacteria interaction but they are costly and time consuming [11-15].

Thus, there is a need to develop computational methods which can predict the correct bacteriophage to treat a bacterial strain. In other words, there is a need to develop a method that can predict phage-host interaction (bacteriophage-bacteria) with high precision. In order to address this problem, a large number of methods have been developed in the past. Broadly, these methods can be classified into three categories as alignment-based, alignment-free and hybrid methods. Following are brief description of major techniques developed in the past for predicting host-phage interaction. WIsH uses alignment free approach to predict prokaryotic hosts of phages using their genomic sequences [16]. VirHostMatcher-Net [17] is hybrid method that combine several alignment-free and alignment-based features. SpacePHARER [18] and VirSorter [19] uses CRISPRs for predicting virus-host interaction in the prokaryotic genomes. PredPHI [20] utilizes virus-host protein-based features for predicting phage-host interactions using deep convolutional networks. These existing methods have their own advantages and disadvantages. Even though number of methods have been developed in the past decade for predicting phage-host interaction yet their accuracy is far from satisfactory. There is a challenge to develop methods that can predict phage-host interaction with high accuracy.

In order to complement existing methods, an attempt has been made to develop an ensemble method for predicting phage-host interaction. This ensemble method combines alignment free and alignment-based methods to improve the accuracy of prediction. In order to maintain standards and compare with existing methods, we developed and evaluate our models on benchmark datasets used in recent study [17]. There were few issues with this benchmark dataset which we removed in current study. We successful demonstrated that the models developed in this study are more accurate than previous methods in predicting phage-host interaction at all levels (e.g., Genus, Family, Phylum). One of the major objectives of this study is to facilitate researchers working in the field of phage therapy. Thus, we developed PhageTB (Webserver and Standalone Software) that contain three major modules; i) host for a phage, ii) phage-host interaction and iii) phage for a host. Our module host for a phage allow users to predict name for bacterial strain (host) from sequence of a virus/phage. Phagehost interaction module allow one to predict whether a given virus and bacterial strain will interact or not. Third module, phage for a host allow user to predict most appropriate phage/virus that can lyse a given strain of bacteria.

## Materials and methods

### Dataset Collection & Pre-processing

In the present study, datasets used for training and validation were obtained from a recent study VirHostMatcher-Net [17]. It contains 25899 distinct species, 2814 different genera, 502 distinct families, 219 distinct orders, 95 distinct classes, 55 distinct phyla, and 2 distinct domains. The training dataset comprises 826 phages and their corresponding hosts (till the strain level), out of which 817 infect bacteria while nine infect archaea. The remaining 1462 phage entries and their corresponding hosts are used as the testing dataset to evaluate and compare the proposed methods’ performance. There are few issues in testing dataset, it contain phages with putative hosts whose lower taxa levels are not present in the training set. Evaluation of such phage-host interaction is not practically feasible as well as advisable. Ideally, test dataset should only contain phage-host interaction whose complete information is available. Hence, we modified the testing dataset by filtering out the phages whose hosts are not represented in the training dataset at the genus level, which lead to 1201 phage-host interactions. Finally, our training dataset have 826 phage-host interactions and testing dataset have 1201 phage-host interaction. In order to make unbiased comparison with existing methods, we also compute the performance on original test dataset that contain 1462 phagehost interactions (See Supplementary Materials). Figure 1 highlights the distinct hosts that are represented in at least one interaction in the train, test, and the modified test sets across seven taxonomic levels, namely Genus, Family, Order, Class, and Phylum. The distribution of

**Figure 1:**
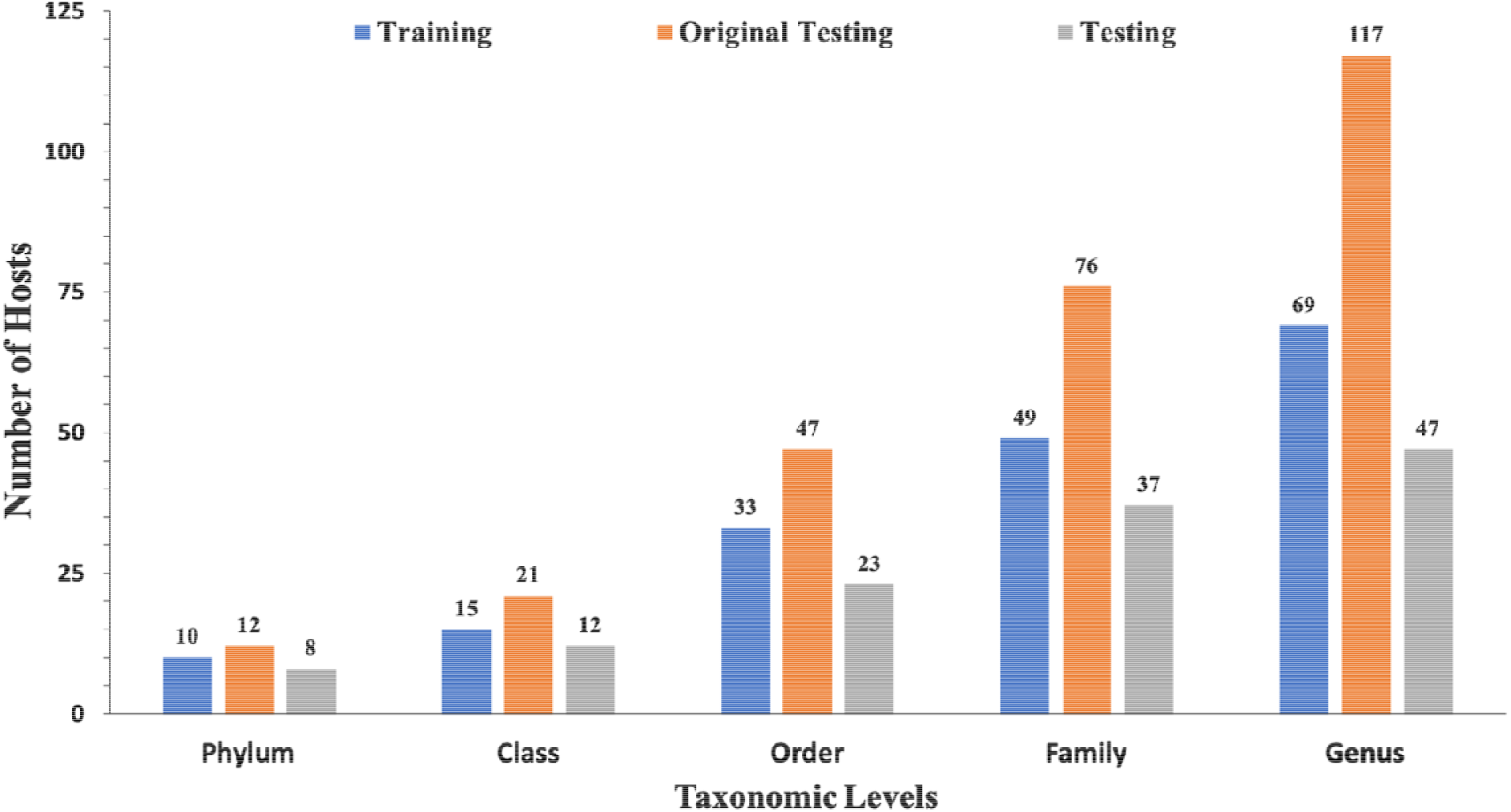
Distribution of data in training, testing, and original testing datasets at different taxonomic levels

### Outline of the study

In this study, we have developed number of alignment-based methods mainly using BLAST called BLAST_Host_, BLASTPhage and CRISPRPred (See Fig 2C). All these alignment-based methods are based on top hits of BLAST search. In addition, models have developed using machine learning techniques where sequence are represented by fixed length vectors. We also developed a hybrid method that combines a machine-learning based model with similarity scores (See Figure 2B). Finally, an ensemble method has been developed that combines all alignment-based models with hybrid method in a sequential method (Fig 2A). The ensemble model employs a sequential prediction process, wherein predictions from BLAST_Phage_ are used if the e-value of alignment is within a predetermined threshold. For the remaining phages, predictions from BLAST_Host_ are used if the e-value is within the predetermined threshold. Predictions from the hybrid model (if the prediction score is above a threshold) are used for the phages whose host could not be predicted from the first two methods. Lastly, for the remaining phages, predictions are made using CRISPR_Pred_.

**Figure 2.**
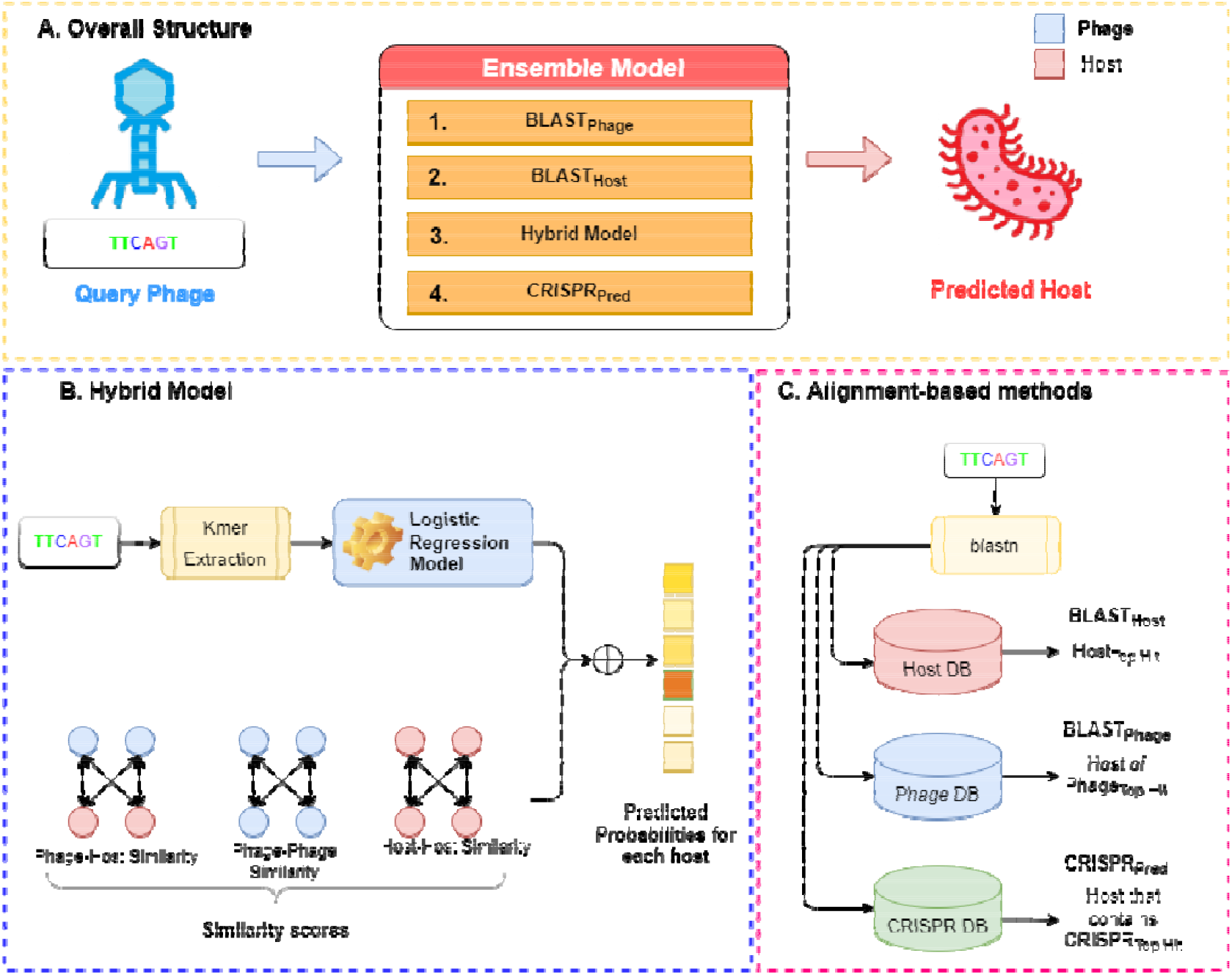
Overall structure of algorithms used in this study; (A) Ensemble method, (B) Hybrid model (C) Alignment-based method

### Alignment-based Methods

Most of the alignment-based methods exploit sequence similarity between genomes of phages and their hosts. The most widely used method for searching similar sequences is BLAST [21]. We employed BLAST-based predictions at three levels, such as BLAST_Phage_, BLAST_Host_, and CRISPR_Pred_ model. In case of BLAST_Phage,_ sequence of a query phage/virus is search against database of phages whose interacting host is already known. In our study, the customized database referred as reference phage database was created using training dataset comprising the information about the phages and their respective interacting hosts. Then, the phage sequences in the testing dataset were searched against the reference phage database using BLAST at different e-values. The host corresponding to the top BLAST hit of a phage is assigned as predicted host for query phage. In summary, BLAST_Phage_ model predict host based on similarity in query and target phage. In case of BLAST_host,_ sequence of a phage is searched in database of 185 host sequences used in training dataset. The top hit from this alignment task was assigned as the potential host. CRISPR systems play a vital role during the infection process of phages and infection prevention by hosts. As a prevention strategy, prokaryotes place a fragment of the genome of an infecting phage as a spacer in the CRISPR array, which is a recognizable repeat region in the genome. Such a sequence indicates a recent infection and thus can be used as a potential signal for predicting hosts. CRISPR Recognition Tool (CRT) [22] is used to identify CRISPR locus in the bacterial genomes using a reference host database. We extracted CRISPR sequences using the CRT tool and created a reference CRISPR database. The test dataset genomes were aligned with the reference CRISPR database using BLAST, where the host corresponding to the top hit is predicted as the potential host. In case of CRISPR alignment we have utilized BLAST short-task parameter as used in previous study [23]. We termed this approach of alignment as CRISPR_Pred_ model.

### Generation of Features

It is important to compute fixed number of features corresponding to each phase for developing a machine-learning based model. These phage sequences are polymer of four nucleotides (A, T, G, C) and have wide range of variation in length. One of the commonly used technique to generate fixed number of features in as sequence is to calculate frequency of words in a sequence. For example one can calculate frequency of nucleotides in a sequence, where sequence can be represented by a vector of dimension four. In this case, total number of words are 4^k^, where world length or k-mer (k) is equal to one. Similarly, frequency of di-nucleotides (i.e, AA, AC, AG, AT, CA, CC) can be calculated, where total number of worlds will be 16 (4^2^) with world length two. One of the limitation of this frequency based features that they are biased with length of sequence and noise in sequence. Thus, in this study we used modified frequency words, which subtract of frequency of world by chance in sequence [24]. Following formula was used to compute modified frequency of words, which is used

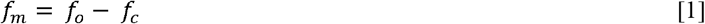

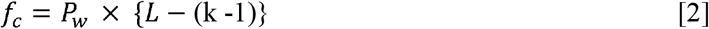

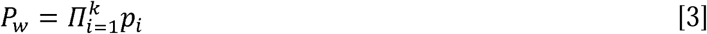

where *f*_*m*_, *f*_*0*_ and *f*_*c*_ are modified, original and chance frequency of a word w respectively. *P*_*w*_ is probability of word whereas *p*_*i*_ is probability of a nucleotide *i* in word *w. L* is length of sequence or number of nucleotides in sequence. K-mer *k* is size or length of word of *w*.

### Machine Learning Model

For this level of classification, several machine learning classifiers were implemented and compared to develop the best performing model. We have implemented various machine learning techniques include Random Forest (RF), Gaussian Naive Bayes (GNB), Logistic regression (LR), Support vector machine (SVM) with a linear kernel, eXtreme Gradient Boosting (XGBoost), Decision Tree (DT), K-Nearest Neighbour (KNN), and Multi-layer Perceptron (MLP). These classification techniques were implemented using the python-library scikit-learn [25].

### Hybrid Model

We have developed various machine learning model at the third level for the remaining phages i.e., the phages whose host could not be predicted using BLAST_Phage_ and BLAST_Host_ method. We term this level of prediction as the hybrid model. Due to the coevolution of phages and their hosts, their genetic compositions are highly similar, thus a given phage has significant overlap with its putative host at the genomic level. Therefore, similar hosts will likely be infected by the same phage, or similar phages are likely to infect the same host. We have used the base machine-learning model prediction probabilities *Pr*_*b*_ *=* [*Pr*^*i*^ *for i =* 1, … *M*] for all hosts, where M is the total number of hosts in the reference host database and *Pr*^*i*^ (prediction probability of i^th^ bacteria) which varies between 0 to 1. In addition, we have added the similarity-scores (SIM) i.e., bit-scores from BLAST alignment tasks between phage-phage, phage-host, and host-host databases using a weighted sum. Further, we have used the Pr_o_ to calculate the final prediction probabilities for all bacterial hosts.

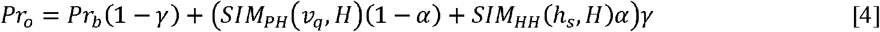

Where, *SJM*_*PP*_, *SJM*_*PH*_ *and SJM*_*HH*_ denote phage-phage, phage-host, and host-host similarity scores, where *SJM*_*PH*_ *(v, H)* gives an M-Dimensional vector that gives the similarity scores of phage v with all hosts in the set *H= [h*_*1*_, *h*_*2*_,*… h*_*M*_] of reference hosts. Similarly, *SJM*_*HH*_ *(h, H)* also gives an M-Dimensional vector denoting the similarity of the host *h* with all other hosts in set H. *v*_*q*_ corresponds to the input query phage, *h*_*s*_ represents the host of most similar phage in the training dataset based on *SJM*_*PP*_, and *Pr*_*b*_ is the prediction probabilities from the base model. Here, α and γ are the weighting parameters used in the given equation and are determined experimentally during cross-validation using grid search over the value range of 0 to 1 with step size of 0.1. The final predictions from the hybrid model were calculated using equation 5.

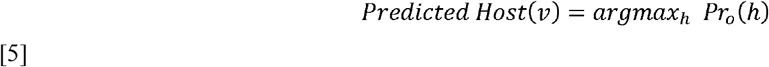

### Ensemble Model

In order to improve the prediction accuracy, without compromising the coverage we have used an ensembled approach, where we have generated predictions using combinations of different models. Here, we tried to integrate alignment-based, alignment-free and hybrid models. At first, we calculate predictions from BLAST_Phage_ and assign host against phages where the e-value of alignment is within a threshold. Similarly, this process was repeated for remining phages using BLAST_Host_. Secondly, we computed predictions from the hybrid model for phages where final prediction score is above a threshold (See Equation 5). Finally, for the remaining phages, predictions are made using CRISPR_Pred_.

### Evaluation Parameters

We evaluate the performance of our approach on the original testing dataset, which comprises 1462 phage samples. Moreover, we have also evaluated the performances of the generated models on the modified testing dataset containing 1201 phage samples. We also compare our approach with past studies in terms of prediction accuracies for correctly predicting hosts binned by taxonomic levels from Genus to Phylum. The prediction accuracy is defined as the fraction of phages whose hosts were identified correctly out of the total phages at a given taxonomic level.

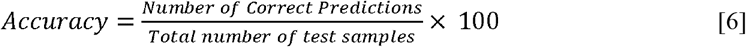

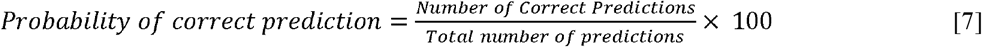

### Webserver Architecture

A web server named as ‘PhageTB’ (https://webs.iiitd.edu.in/raghava/phagetb/) is developed to predict the bacterial hosts, host-phage interactions and lytic phage for a bacteria. The front end of the web server was developed by using HTML5, JAVA, CSS3 and PHP scripts. It is based on responsive templates which adjust the screen based on the size of the device. It is compatible with almost all modern devices such as mobile, tablet, iMac and desktop.

## Results

### Predictions from BLAST_Phage &_ BLAST_Host_

Sequence alignment of phage and host genomes is the primary method for assigning hosts to phages from a set of known hosts. For this purpose, we employed BLAST technique, where first we vary the degree of alignment by changing the threshold on e-value. We observed that the similarity search improved the accuracy. When the e-value threshold is reduced, it leads to more specific aligned hits and thus improved prediction accuracies for the samples we get a hit for, but the overall recall decreases. Whereas for many phages, we cannot predict hosts using this method as shown in Figure 3. When aligning the phage genomes in the original testing dataset with the reference phage genome database, and assigned the host based on top hit, we attained the prediction accuracies of 45.2%, 56.2%, 62.8%, 67.5%, and 71.2% at Genus, Family, Order, Class and Phylum levels, respectively (Supplementary Table S1).

**Figure 3.**
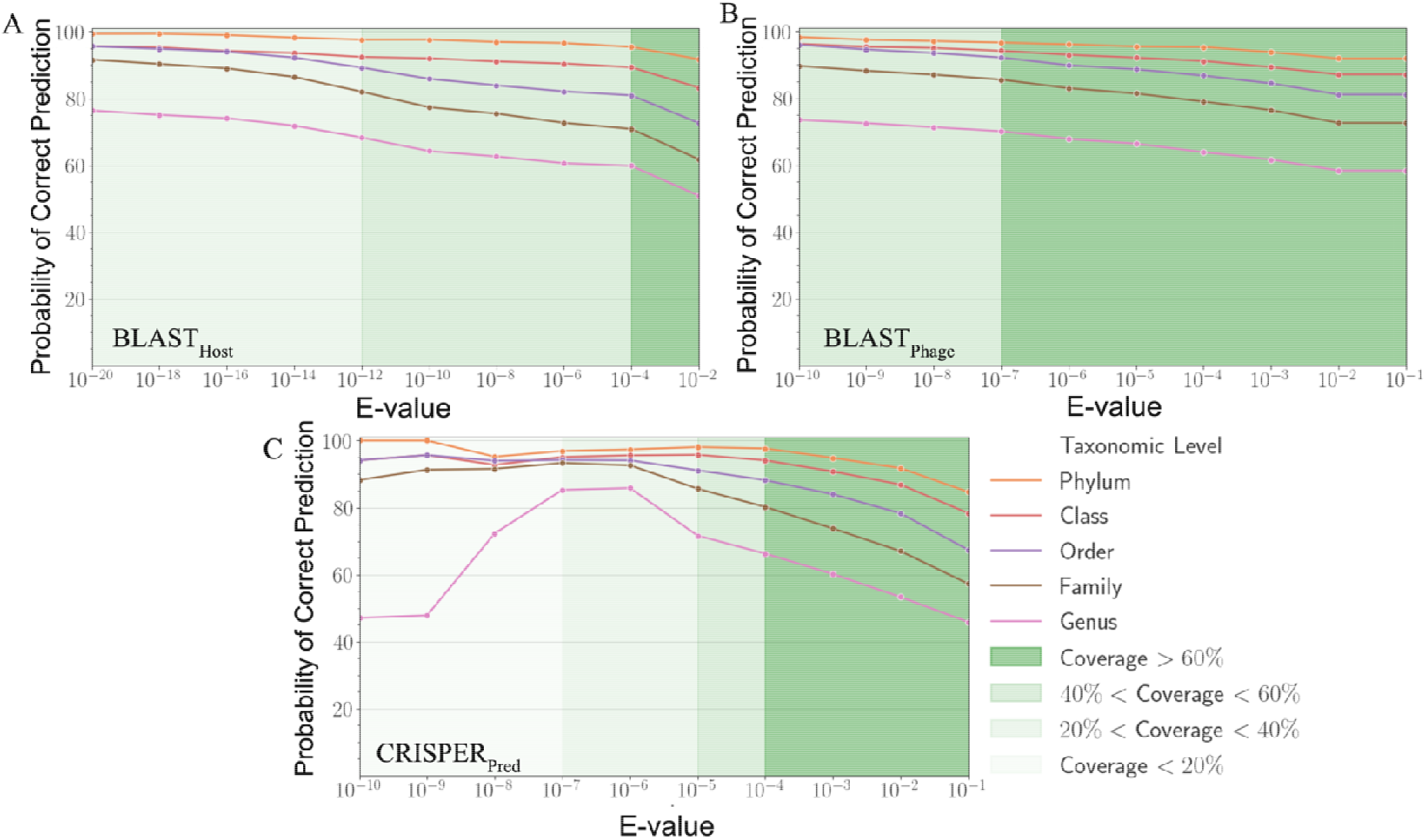
Variation in probability of correct prediction (A) BLAST_Phage_, (B) BLAST_Host_ and (C) CRISPER_Pred_ at different e-values

Similarly, on aligning the phage genomes with reference host genome database and assigning the top hit as the predicted host, we obtained the accuracies of 34.8%, 42.3%, 49.7%, 57.0%, and 62.8% at Genus, Family, Order, Class and Phylum levels, at e-value 1.00E-02 (Supplementary Table S1). As reported in Table 1, we obtained accuracies of 42.4%, 50.5%, 57.2%, 61.4%, and 66.2% at different taxonomic level using BLAST_Host_ method at e-value 1.00E-02. Similarly, BLAST_Phage_ attained accuracies of 55.0%, 66.4%, 71.4%, 74.9%, and 78.4% at Genus, Family, Order, Class and Phylum levels, respectively on the test dataset. Further predictions were made via aligning phage genomes with CRISPR sequences extracted from host genomes. We observed that in case of original test dataset (See Supplementary Table S1) and modified test datasets, the prediction accuracies were improved at the level of class and phylum in comparison with BLAST_Host_ and BLAST_Phage_ methods (refer to Table 1).

**Table 1:**
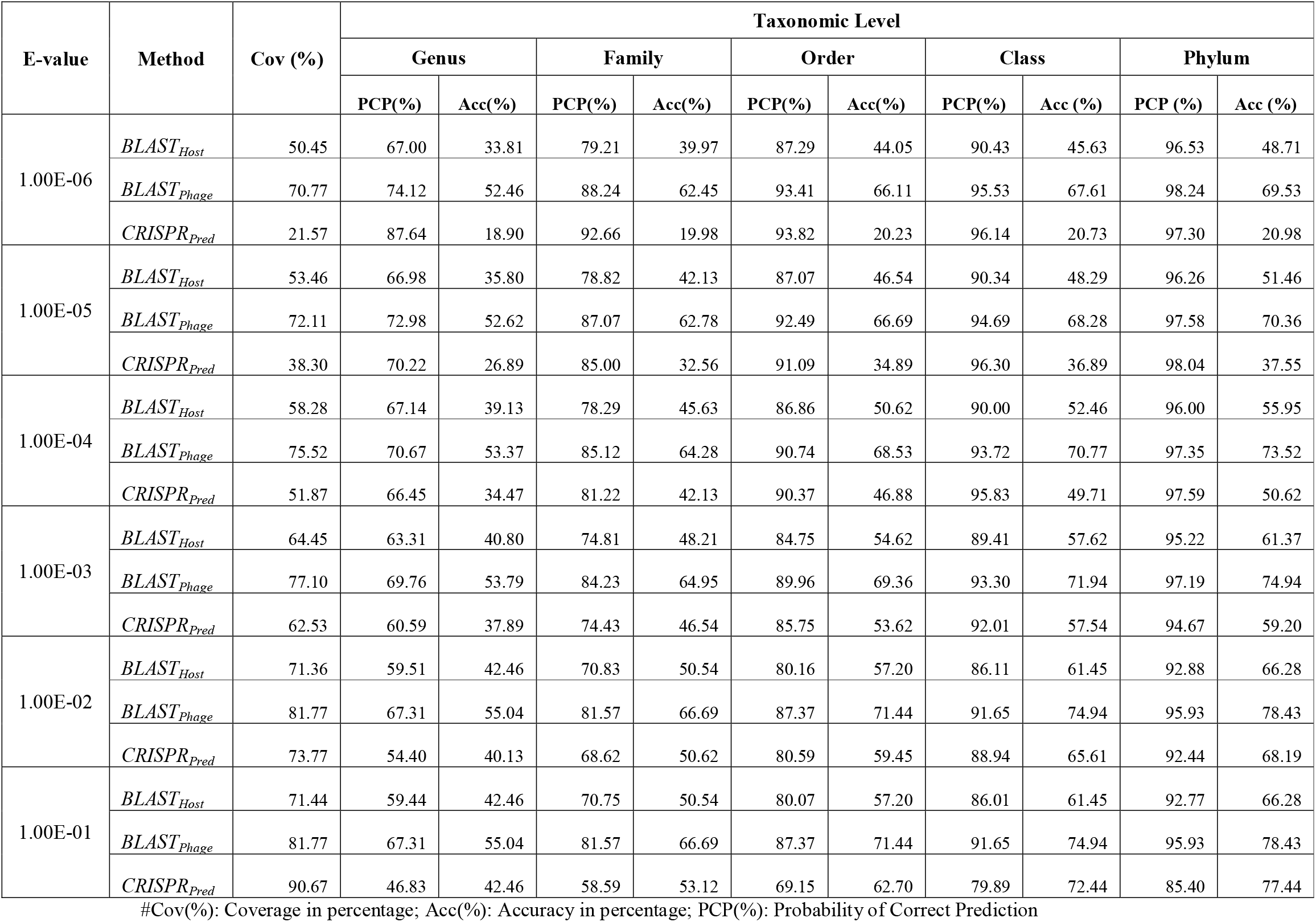
Prediction of five taxonomic levels of bacterial host using alignment-based models on validation dataset.

### Performance of Machine learning models

In order to develop various machine learning models i.e., Decision Tree (DT), Random Forest (RF), Gaussian Naive Bayes (GNB), XGBoost, Logistic Regression (LR), Multi-layer perceptron (MLP), and Support Vector Machine (SVM), we extracted the features using *f*_*m*_ equation 1 with k = 6, from the phage genomes and predicts the host class on validation dataset. As represented in Table 2, on modified test dataset, LR-based models performed best among all other classifiers. In order to improve the performance we integrated the prediction score of the best model i.e., LR with the similarly score i.e., BLAST and observed that there is a significant improvement in the predictive accuracies. Further, we have trained the hybrid model by varying the weighting parameters α and γ in equation 4; we achieved maximum performance at α = 0.9 and γ = 0.6. On original test dataset, the prediction accuracy of hybrid model (LR + similarly score) is 49.7%, 64.7%, 75.3%, 84.8%, and 90.6% at different taxonomic levels (Supplementary Table S2). On the other side, the hybrid model developed using modified test dataset, outperformed all other classifiers with an improved accuracies of 60.6%, 75.8%, 82.0%, 89.7%, 93.5% at Genus, Family, Order, Class and Phylum levels (See Table2).

**Table 2:** Prediction of five taxonomic levels of bacterial host using machine learning and Hybrid models on modified test dataset.

| Models | Taxonomic Levels (Accuracy %) |  |  |  |  |
| --- | --- | --- | --- | --- | --- |
|  | Genus | Family | Order | Class | Phylum |
| Decision Tree (DT) | 22.30 | 30.50 | 34.70 | 45.70 | 55.90 |
| Gaussian Naive Bayes (GNB) | 24.30 | 32.00 | 35.20 | 42.90 | 45.60 |
| XGBoost (XGB) | 37.20 | 44.20 | 48.20 | 53.20 | 57.80 |
| Random Forest Classifier (RF) | 38.80 | 46.10 | 51.00 | 55.70 | 59.20 |
| Linear SVM (SVM) | 51.80 | 62.10 | 65.60 | 71.00 | 73.50 |
| K-Nearest Neighbor (KNN) | 49.20 | 61.60 | 67.60 | 73.20 | 78.50 |
| Multi-layer perceptron (MLP) | 49.20 | 63.50 | 69.10 | 76.10 | 80.00 |
| Logistic Regression (LR) | 54.80 | 68.10 | 72.50 | 77.70 | 80.80 |
| Hybrid Model (Similarly-scores + LR) | 60.60 | 75.80 | 82.00 | 89.70 | 93.50 |

**Table 3:** Prediction of five taxonomic levels of bacterial host using ensembled models.

| Method | Taxonomic Level (Accuracy %) |  |  |  |  |
| --- | --- | --- | --- | --- | --- |
|  | Genus | Family | Order | Class | Phylum |
| BLAST<sub>Phage</sub> + BLAST<sub>Host</sub> | 59.40 | 70.10 | 73.90 | 75.50 | 78.60 |
| BLAST<sub>Host</sub> + Hybrid Model | 65.00 | 78.10 | 83.10 | 89.90 | 93.90 |
| BLAST<sub>Phage</sub> + Hybrid Model | 62.60 | 75.80 | 82.40 | 89.80 | 93.60 |
| CRISPR<sub>Pred</sub> + Hybrid Model | 61.10 | 74.10 | 80.50 | 86.40 | 90.10 |
| BLAST<sub>Phage</sub> + BLAST<sub>Host</sub> + Hybrid Model | 65.70 | 78.60 | 84.30 | 90.70 | 94.00 |
| BLAST<sub>Host</sub> + CRISPR<sub>Pred</sub> + Hybrid Model | 65.50 | 76.60 | 81.70 | 86.80 | 90.90 |
| BLAST<sub>Phage</sub> + CRISPR<sub>Pred</sub> + Hybrid Model | 66.70 | 79.20 | 84.60 | 89.80 | 93.50 |
| BLAST<sub>Phage</sub> + BLAST<sub>Host</sub> + CRISPR<sub>Pred</sub> + Hybrid Model | 67.90 | 80.60 | 85.50 | 90.00 | 93.50 |

### Performance of Ensemble Models

In order to improve the performance of above mentioned models, we have used ensembled approach, where we have generate predictions on the combinations of different models. Here, we have tried each and every combinations of BLAST_Phage_, BLAST_Host_, CRISPR_Pred_ and Hybrid Model and validate the accuracies at different taxonomic levels on both testing data. We observe improvements across taxonomic levels as we progressively add different prediction methods to the overall framework. Host prediction accuracies were markedly higher than individual components. For higher-order taxonomic levels (Class and Phylum) combination of BLAST and hybrid model based predictions also got comparable results, however for lower levels, the best performing approach was the one that combines all prediction methods. Our proposed ensembled model (BLAST_Phage_ + BLAST_Host_ + CRISPR_Pred_ + Hybrid Model) outperforms the existing approaches across all taxonomic levels, correctly predicting 61.6%, 74.4%, 80.5%, 85.7%, and 91.2%, respectively, for original test dataset (Supplementary Table S3) and 67.9%, 80.6%, 85.5%, 90.0%, and 93.5% for test dataset at Genus, Family, Order, Class and Phylum levels. The e-value thresholds for BLAST_Phage_ (1.00E-10), BLAST_Host_ (1.00E-20), CRISPR_Pred_ (1.00E-2), and the prediction probability threshold for the hybrid models is 0.6.

### Comparison with other Methods

It is very important, to compare this newly developed method with the existing tools to understand the merits and demerits. Therefore, we compare the performance of our method with three existing tools (PHP [26], VirHostMatcher-Net [17], and Phirbo [27]), as shown in Figure 4. As shown below PhageTB outperform previous studies at each taxonomic level, with an accuracy of 67.90%, 80.60%, 85.5%, 90.0%, and 93.5% at Genus, Family, Order, Class and Phylum levels. The prediction accuracies of other tools is provided in Figure 4.

**Figure 4.**
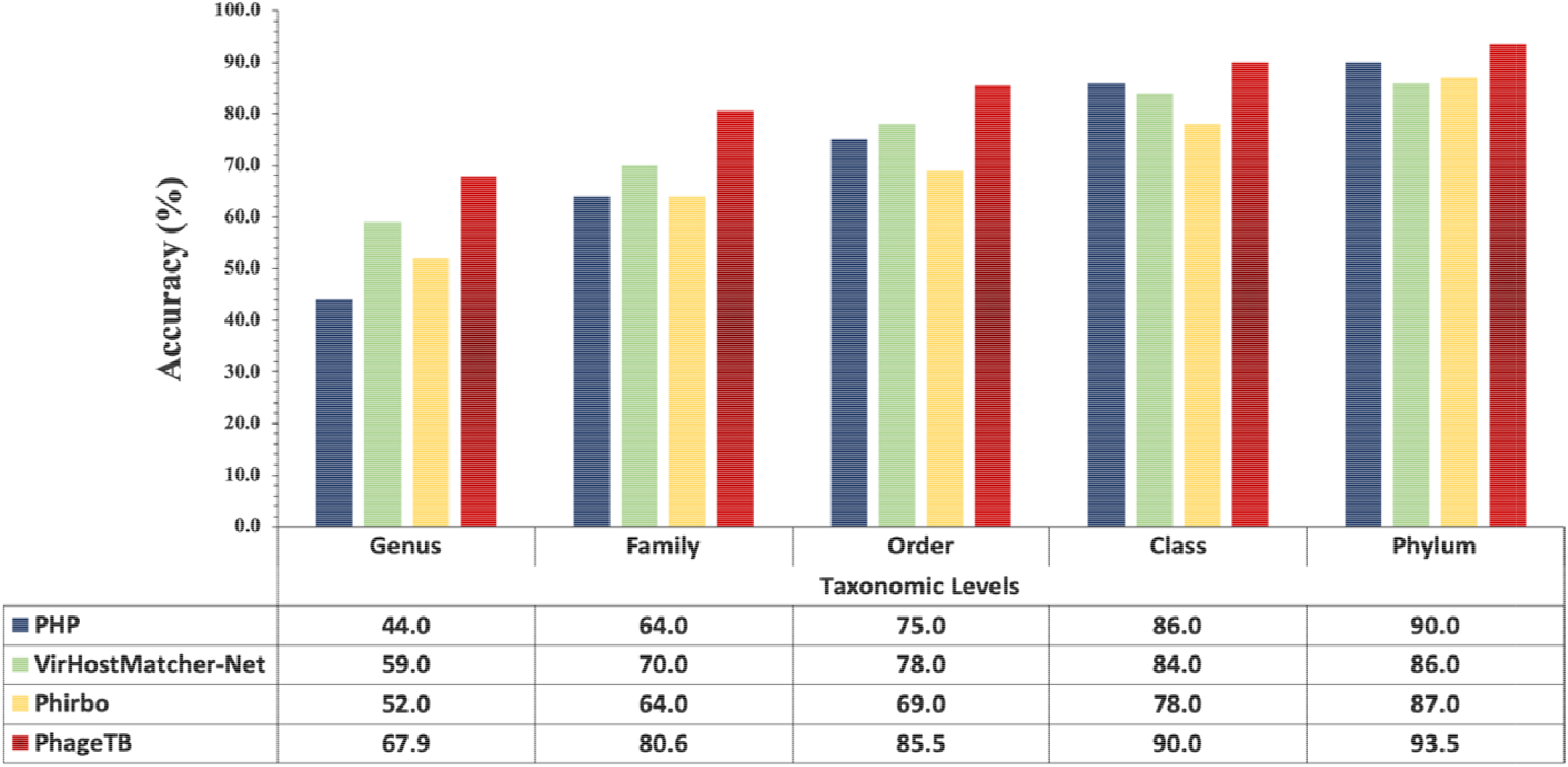
Comparison of performance of our method with existing tools at different taxonomic levels

### Contributions to the Scientific Community

For serving the scientific community, we integrate our best models in a webserver named “PhageTB”. This tool incorporates three major modules (i) Hosts for bacteriophages (ii) Interaction of phage-host pair and (iii) Lytic phage for a bacterial host. The first module “Hosts for bacteriophages” allows a user to choose four predictive methods i.e., BLAST_Phage_, BLAST_Host_, CRISPR_Pred_ and Hybrid Model. User need to provide the query genome sequence and our tool predict the bacterial hosts using the reference host database. The second module “Interaction of phage-host pair” predicts whether a given pair of phage and bacteria are likely to interact based on their genome sequences. User need to provide genome sequences of phage and bacteria in the FASTA format and our tool predict the interactions between the query sequences and the predictions were entirely based on the alignment score. User can choose any prediction method and our tool can predict the interactions based on the similarity-based alignment, i.e., BLAST. The third module “Lytic phage for a bacterial host” predicts bacteriophages corresponding to query bacterial sequences. The input genome sequence are searched against a reference database of phage-host interactions, where first we align the query sequence with genome sequences of bacteria that are known hosts for some bacteriophages. The top hit bacteria from the reference database are the most similar bacteria to the query, and thus the query is likely to be infected by the phage associated with the top hit.

The webserver “PhageTB” was implemented using HTML, CSS, and PHP and has multidevice compatibility, and provides an easy-to-use and user-friendly interface. The opensource web server is available at https://webs.iiitd.edu.in/raghava/phagetb. The command line standalone can be found on GitHub at https://github.com/raghavagps/phagetb.

### Case study: Prediction of Lytic Phages

Predicting lytic phages that can be used (solely or with other agents) for treatment of multidrug resistance bacterial infections is a major problem of concern for the scientific community [6, 28]. In this case study we identify suitable phage-based treatments for drugresistant bacterial infections and we use our webserver PhageTB to predict the lytic phages corresponding to the six ESKAPE (Enterococcus faecium, Staphylococcus aureus, Klebsiella pneumoniae, Acinetobacter baumannii, Pseudomonas aeruginosa, and Enterobacter species [29, 30] bacteria. It comprise the six well known highly virulent antibiotic-resistant bacterial pathogens. Here we have download the genome assemblies of each of the six bacteria from NCBI (https://www.ncbi.nlm.nih.gov/assembly/) and predict the specific phage. We utilize the default parameters of PhageTB “Lytic phage for a bacterial host” module for the prediction of phages that are likely to infect a bacterium. Table 4 and Supplementary Table S4 represents the predicted phages with GenBank ID for five out of six ESKAPE bacteria. We were not able to predict any lytic phage against Pseudomonas aeruginosa bacteria, however we evaluated the predictions of our tool with existing studies and clinical trials [29, 31] (Refer Supplementary material). These findings can be extended to other drug-resistant bacterial strains and thus utilized to expedite the process of finding suitable phages for the treatment of drug-resistant bacterial infections where the lytic phages are not known beforehand.

**Table 4:**
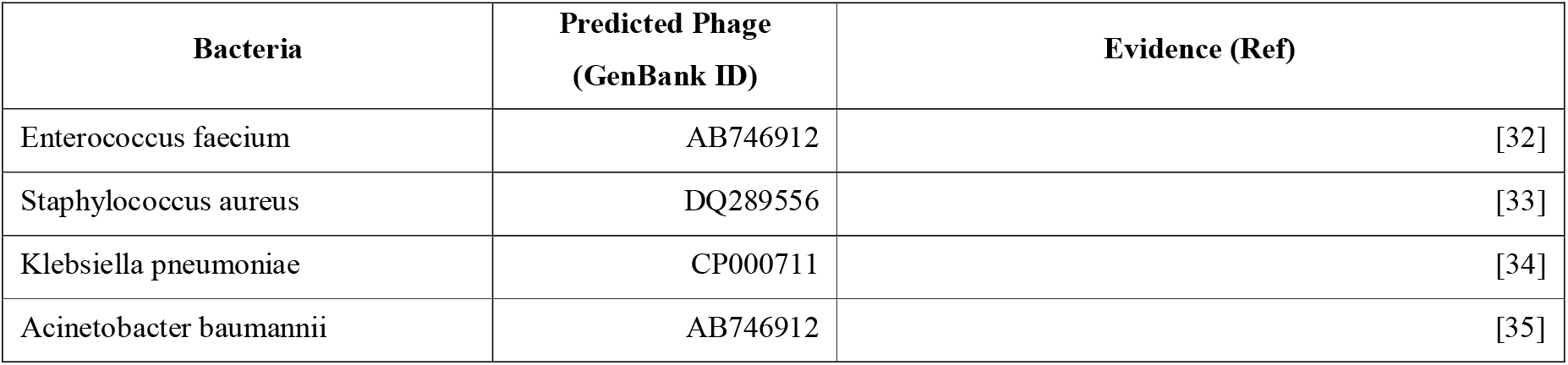
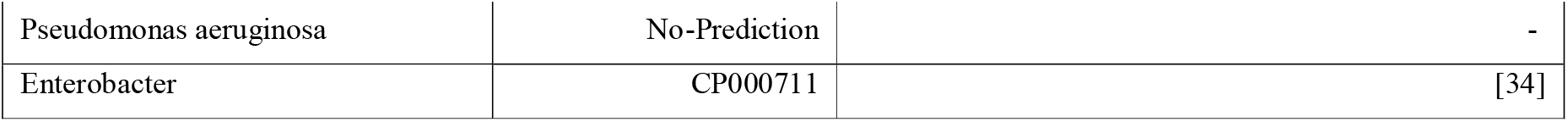
Lytic phage prediction by phageTB on ESKAPE bacteria.

| Bacteria | Predicted Phage<br>(GenBank ID) | Evidence (Ref) |
| --- | --- | --- |
| Enterococcus faecium | AB746912 | [32] |
| Staphylococcus aureus | DQ289556 | [33] |
| Klebsiella pneumoniae | CP000711 | [34] |
| Acinetobacter baumannii | AB746912 | [35] |
| Pseudomonas aeruginosa | No-Prediction | - |
| Enterobacter | CP000711 | [34] |

## Discussion and Conclusion

Phage therapy is a leading alternative to antibiotics for the treatment of bacterial infections as most pathogenic strains are now showing resistance to numerous known antibiotics [36]. The development of phage therapy requires the identification and isolation of a large number of bacteriophages. Phages are generally specific to bacterial species as well as their strains which is an advantage of this therapy as it will only kill the pathogenic bacteria, leaving out the natural bacteria required for the human body. The highly specific nature of bacteriophages necessitates the collection and characterization of their known and potential hosts as well as the interactions between them [37, 38]. There have been several studies in the past that have made an effort to identify and predict the hosts of phages and their interactions like WIsH, VirHostMatcher-Net, SpacePHARER, VirSorter, and PredPHI [16-20]. In spite of this, the presently available methods cannot accurately predict the taxonomic classes of the phage and hosts. To bridge this gap and achieve higher accuracies in predicting the phagehost interactions, we developed a method called PhageTB that uses both alignment-based and alignment-free features to predict the hosts from query genomic sequences of bacteriophages.

PhageTB is a hierarchical prediction method that stacks four predictive methods to predict the phage-host interactions across five levels – Genus, Family, Order, Class, and Phylum.

These methods include BLAST_Phage_, BLAST_Host_, Hybrid model, and CRISPR_Pred_.BLAST_Phage_, BLAST_Host_, employ BLAST alignment based predictions for query sequences against reference hosts and phages respectively. The Hybrid Model predicts the host based on the machine learning classifier and similarity scores whereas the CRISPRPred approach uses CRISPR based alignment to predict the same. These four methods combined together are able to accurately predict the host-phage interactions and outperform the previously developed methods to predict phage-host interactions namely PHP, VirHostMatcher-Net, and Phirbo when tested on a dataset containing 1,462 phage-host interactions [17, 26, 27]. We obtained 67.9%, 80.6%, 85.5%, 90.0%, and 93.5% accuracies for Genus, Family, Order, Class, and Phylum respectively using ensemble method which is higher than the accuracies obtained from the abovementioned methods. The various approaches combined together in PhageTB provide with the accurate predictions for phage-host interactions which makes it a valuable tool for the scientific community working in this field worldwide to combat the crisis of antibiotic resistance. With the increasing availability of metagenome samples, new methods for identifying phages and determining their hosts are required. We believe that PhageTB will prove to be an effective tool in finding specific hosts for the phages which can be potentially helpful in the development of phage therapy, ecology research, viral metagenomics analysis, and human gut microbiocenosis research along others. PhageTB is an easy-to-use method of assigning hosts to bacteriophages, studying their interactions, and narrowing down the search space for candidate phages that can successfully lyse the query bacteria and thus be utilized in phage therapy for treating bacterial infections caused by it. Our tool is freely accessible at https://webs.iiitd.edu.in/raghava/phagetb/index.html, python standalone package is available at https://webs.iiitd.edu.in/raghava/phagetb/down.php and GitHub at https://github.com/raghavagps/phagetb.

## Authors’ contributions

SA, and GPSR collected and processed the datasets and implemented the algorithms. SA, AD, and SP created the back-end of the web server and front-end user interface. SA developed the prediction models. SA, SP, SC, and AD analyzed the results. SA, AD, SC, AA, SP, and GPSR penned the manuscript. GPSR conceived and coordinated the project and provided overall supervision to the project. All authors have read and approved the final manuscript.

## Acknowledgements

Authors are thankful to the Department of Bio-Technology (DBT) and Department of Science and Technology (DST-INSPIRE) for fellowships and the financial support and Department of Computational Biology, IIITD New Delhi for infrastructure and facilities.

## Data Availability Statement

All the datasets generated in this study are available at https://webs.iiitd.edu.in/raghava/phagetb/down.php

